# Cancer Classification from Healthy DNA

**DOI:** 10.1101/517839

**Authors:** Siddharth Jain, Bijan Mazaheri, Netanel Raviv, Jehoshua Bruck

## Abstract

The genome is traditionally viewed as a *time-independent* source of information; a paradigm that drives researchers to seek correlations between the presence of certain genes and a patient’s risk of disease. This analysis neglects *genomic temporal changes*, which we believe to be a crucial signal for predicting an individual’s susceptibility to cancer. We hypothesize that each individual’s genome passes through an *evolution channel* (The term *channel* is motivated by the notion of communication channel introduced by Shannon1 in 1948 and started the area of *Information Theory*), that is controlled by hereditary, environmental and stochastic factors. This channel differs among individuals, giving rise to varying predispositions to developing cancer. We introduce the concept of *mutation profiles* that are computed without any comparative analysis, but by analyzing the short tandem repeat regions in a *single healthy genome* and capturing information about the individual’s evolution channel. Using machine learning on data from more than 5,000 TCGA cancer patients, we demonstrate that these *mutation profiles* can accurately distinguish between patients with various types of cancer. For example, the pairwise validation accuracy of the classifier between PAAD (pancreas) patients and GBM (brain) patients is 93%. Our results show that healthy unaffected cells still contain a cancer-specific signal, which opens the possibility of cancer prediction from a healthy genome.

## Main

The human genome has evolved over time by an interplay of mutational events. An enhanced understanding of the genome’s evolution has numerous direct and practical applications in improving healthcare, discerning ancestry, materializing DNA storage and designing synthetic biology devices for computation. Traditionally, the genome has been viewed as a *time-independent source* of information, and hence much of the genomic research has been focused on discovering variants that cause a certain phenotype. Linkage studies have discovered genes for *Mendelian* diseases such as Cystic Fibrosis^2^, Huntington disease^3^, Fragile-X syndrome^4^ and many others^5^ by investigating genetic variants across families. For more complex diseases, Genome Wide Association Studies (GWAS)^6^ can be used to discover large amounts of risk factors working in conjunction. The broader scope of GWAS has led to the discovery of several new genes and pathways^7^, but many diseases still remain unexplained. Instead of searching for disease-causing variants, we view the genome as a *time-dependent signal*, searching for indicators for how the genome is mutating over time. This gives rise to the following question - What are the possible ways to measure the evolution of mutations? Put differently, how can we quantify the accumulation of mutations in the genome of an individual?

Our approach for extracting time-dependent information about a person’s mutation history is to focus on the tandem repeat regions of this person’s *healthy* genome. We have studied two types of mutations in the genome: tandem duplications and point mutations. Tandem duplications involve the consecutive repetition of a subsequence (e.g. *T<u>CAT</u>G* → *T<u>CATCAT</u>G*). Point mutations, which include substitutions, insertions, and deletions, are single changes in the DNA (e.g. *AC<u>T</u>G* → *AC<u>A</u>G*). When these two processes occur in the same location, point mutations can propagate through tandem duplications, leaving a change in the repeated sequence (see Figure 1a and Methods section). This allows us to construct a likely history of tandem duplications and point mutations. Slippage events can cause regions with many tandem duplications^8^, which are a convenient locations to observe this interaction between mutation processes. In a sense, these *tandem repeats regions* are a nature given *repetition error-detecting code*^9^, where the point mutation errors in the copies store information about the history of the evolution of these regions. These repeat regions effectively characterize an *evolution channel*, which can shed light on the accumulation of mutations in the genome.

**Figure 1.**
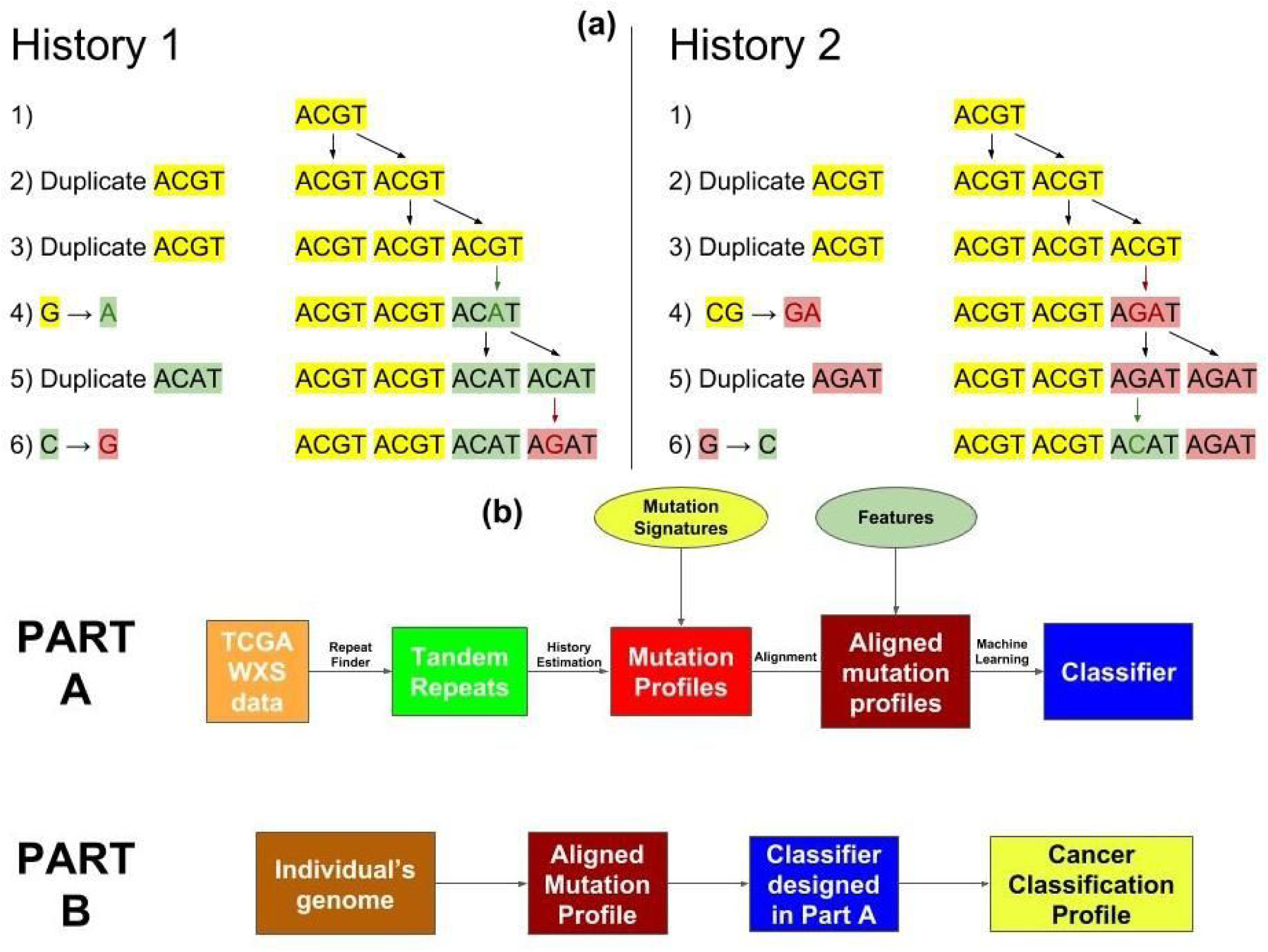
**(a)** Two different evolution histories for the tandem repeat region *ACGTACGTACATAGAT* with pattern length 4 and repeat region length 16. In History 1, only 2 point mutations were needed (marked with green and red respectively). In History 2, 3 point mutations were needed: 1 marked with green and 2 marked with red. In our approach, we would consider History 1 to be more likely as it involves lesser number of point mutations. Therefore, for this tandem repeat region we have that *m* = 2 and *d* = 4. **(b)** The workflow of our algorithm. In Part A a classifier is trained based on the mutation profiles generated from the healthy DNA of cancerous individuals. This part in only performed once per training set. In Part B, the resulting classifier is applied over a given genome to assess an individual’s inclination of developing different cancers.

### Cancer Genomics

Cancer is currently the second leading cause of death worldwide^10^. Cancer is caused by an intricate mixture of complex factors whose inter-relations are not well understood. While the roles of environmental and hereditary factors are well accepted, recent studies suggest that two-thirds of the mutations in human cancers are caused by replication errors^11^.

Most GWAS studies on cancer risk have focused on differences between healthy (i.e., normal) and tumor DNA samples, namely in Single Nucleotide Polymorphisms (SNPs) and Copy Number Variations (CNVs). These studies have discovered tumor suppressor genes like BRCA1, BRCA2, TP53 and oncogenes like HER2 and RAs family^12^. Previous work has also shown that tumor genomes have significantly more genes with repeat instabilities, linking microsatellite instability to colorectal^13^ and other cancers ^13–18^. Another recent approach identified 21 signatures for mutational processes in human cancer using healthy genome based on 96 substitution classifications that were defined by 6 single base substitution classes and the sequence context left and right of the mutated base^19^.

Unlike previous works, we aim to study cancer risk factors while using a *healthy genome*, without using the genome of the tumor itself, which opens the door to cancer prediction and risk assessment. To do this, we analyzed tandem repeat region data in different cancer types from The Cancer Genome Atlas (TCGA)^20^. We estimated the number of point mutations (*m*) and tandem duplications (*d*) in each tandem repeat region by predicting the evolutionary history of those regions^21, 22^ (see Figure 1a). We used the aggregate of this evolution information to form what we call the *mutation profile* of the genome. We then used a gradient boosting algorithm to learn the association between these mutation profiles and the probability of developing specific cancers^23^. The association between the mutation profile and the cancer-type signifies the presence of a cancer-type “signal” in the mutation profiles of the *healthy* genome, which could be useful for future cancer prediction and early cancer detection.

## Results

We hypothesized that different genetic mutation processes accounted for varying risks of developing cancer, and that these processes would leave detectable signals in an individual’s profile. Rigorous verification of this hypothesis would require DNA samples from cancer patients before the onset of their cancer. Such a dataset is not currently available, but blood derived DNA is accessible on The Cancer Genome Atlas (TCGA)^20^, and closely resembles the DNA of cancer patients before they developed the disease. From TCGA, we gathered 3874 *unamplified* blood derived WXS samples which spanned 12 cancers: TCGA-GBM, LUAD, LUSC, PRAD, PAAD, STAD, HNSC, BLCA, KIRC, LGG, SKCM, THCA (Table 1A (Column 2), Supplementary Files 1-12). We used microsatellites (tandem repeats with pattern lengths ≤ 10 bp) with at most 100 repeats to obtain mutation profiles (see Methods, Figure 1).

**Table 1.** **(A)** Number of unamplified healthy samples used for each cancer type in the study showing the number of blood derived normal and solid tissue normal samples. In total, the number of blood derived healthy samples are 3874 and the tissue derived healthy samples are 687. The sample information is provided in Supplementary Files 1-12. **(B)** Number of amplified healthy samples used for each cancer type in the study showing the number of blood derived normal and solid tissue normal samples. In total, the number of blood derived healthy samples are 331 and the tissue derived healthy samples are 194. The sample information is provided in Supplementary Files 13-15.

| **A: Unamplified Samples** |  |  |
| --- | --- | --- |
| Cancer | Blood Derived Normal | Solid Tissue Normal |
| SKCM | 344 | 0 |
| PAAD | 153 | 31 |
| STAD | 396 | 49 |
| BLCA | 393 | 20 |
| PRAD | 440 | 56 |
| LGG | 513 | 0 |
| LUAD | 411 | 102 |
| THCA | 432 | 68 |
| LUSC | 316 | 180 |
| HNSC | 190 | 0 |
| GBM | 255 | 2 |
| KIRC | 31 | 179 |
| **B: Amplified Samples** |  |  |
| Cancer | Blood Derived Normal | Solid Tissue Normal |
| GBM | 171 | 0 |
| LAML | 0 | 135 |
| OV | 160 | 59 |

### Pairwise Cancer Classifiers - Using only Blood Derived Normal Samples

Here we use 3843 unamplified blood-derived normal samples spanning 11 cancers for our analysis (see Table 1A, Supplementary Files 1-12). We did not use the blood derived samples from TCGA-KIRC in this analysis as we only had 31 samples for KIRC which was not enough to construct a reliable classifier. We verified the existence of cancer-type signals within the mutation profiles of blood-derived normal samples by training cancer classifiers using xgboost^23^ and testing their accuracy on separate *validation-set* data (see Methods, Code/Software, Figure 2).

**Figure 2.**
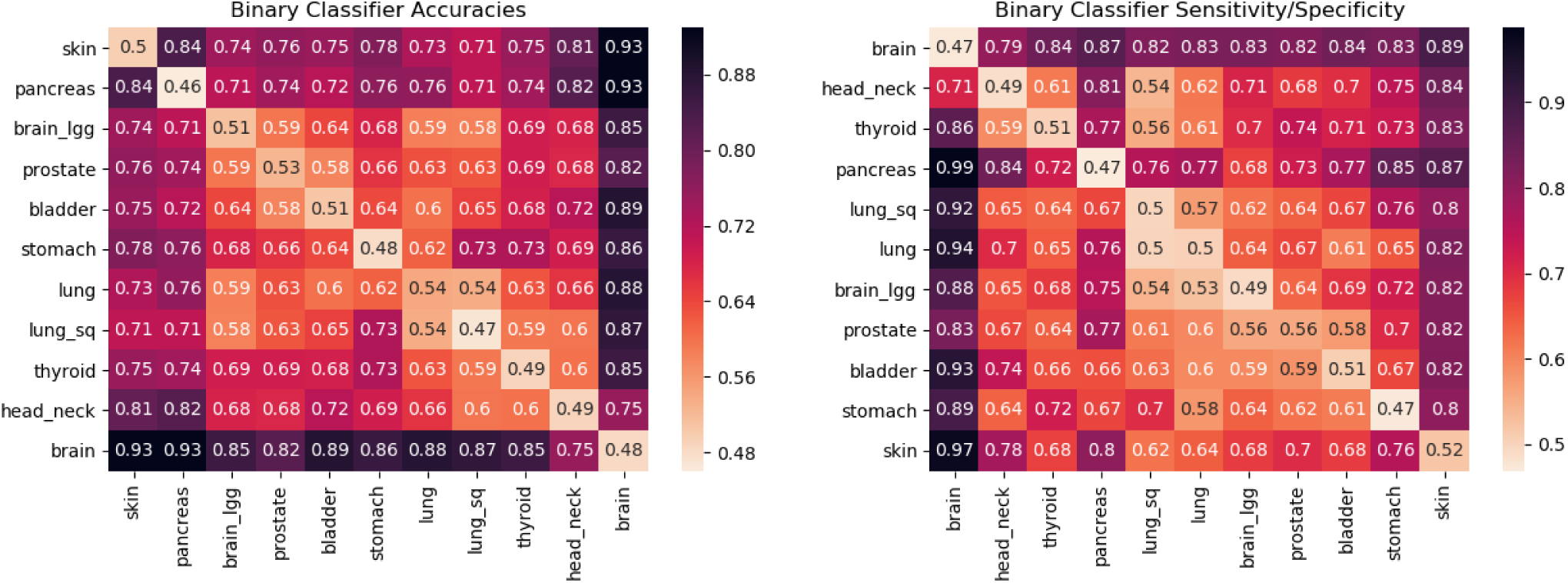
The matrix on the left contains the pairwise validation accuracies of classifiers built by using 3843 blood-derived normal DNA samples covering 11 different cancer types. These cancer types are TCGA-SKCM (skin), PAAD (pancreas), STAD (stomach), BLCA (bladder), PRAD (prostate), LGG (brain_lgg), LUAD (lung), THCA (thyroid), LUSC (lung_sq), HNSC (head_neck), GBM (brain). Each cell in the accuracy seriation matrix represents the average validation accuracy of the binary pairwise classifiers. Each pairwise classifier between cancer X and cancer Y (X≠Y) was constructed using 4-fold cross-validation with patients of each cancer type. These accuracies can be interpreted as distances. The darker the cell, farther are the cancers being compared. The darker rows corresponding to brain, skin and pancreas are indicative of the presence of cancer-type signal in the blood-derived normal (healthy) DNA of cancer patients. The diagonal entries in the seriation matrix represent the accuracies when half of the patients of cancer X were labeled 0 and half of the patients of the same cancer X were labeled 1. As one can expect, the average test accuracy for such classifier should be around 50%. The value in the cell corresponding to the row “pancreas” and the column “prostate” signifies that an average of 74% of the people were correctly classified in each validation pass. The matrix on the right, contains the sensitivity/specificity values. Each cell in the sensitivity/specificity seriation matrix represents the sensitivity value when the row cancer is considered positive and the column cancer is considered negative. It can also be regarded as specificity when the row cancer is considered negative and the column cancer is considered positive. Sensitivity is defined as *TP/*(*TP* + *FN*) and specificity is defined as *TN/*(*TN* + *FP*), where *TP* = True Positive, *FP* = False Positive, *TN* = True Negative, *FN* = False Negative. A value of .77 in the row “prostate” and the column “pancreas” means that 77% of the prostate patients in the test set were truly classified as prostate type (sensitivity when prostate is considered positive). A value of .73 in the row “pancreas” and the column “prostate” means that 73% of the pancreas patients in the test set were truly classified as pancreas type (specificity when prostate is considered positive). The seriation ordering is obtained by solving TSP (i.e., the Travelling Salesman Problem) exhaustively^24^.

As can be seen in Figure 2, mutation profiles of blood-derived normal DNA of GBM patients shows strongly distinctive signals from the rest of the tested cancers with classification accuracies ranging in between 75% for HNSC to as high as 93% for SKCM and PAAD. A similar observation is made for both SKCM and PAAD as they are distinguishable from all of the other cancers with more than 71% accuracy. For other cancers - STAD, BLCA, LGG, PRAD, LUAD, THCA, LUSC, HNSC, the distinguishing signal is much weaker for many cancers. For example, LGG when compared against PRAD, BLCA, LUAD, LUSC gives pairwise accuracies of 59%, 64%, 59% and 58% respectively. Cancers with risk factors that emit different mutation profiles are easier to distinguish, resulting in more accurate classifiers. Hence, accuracy gives a notion of distance on the scale of 50% (close, indistinguishable) to 100% (far, different). The order of the cancers in the display minimizes the distances between neighboring cancers using the travelling salesman problem (TSP)^24^, giving a likely low dimensional projection of the features being learned by the classifiers. The observed accuracies and specificity/sensitivity observed in Figure 2 confirm the presence of *cancer-type* signal in the blood-derived normal DNA.

The clustering of cancers in Figure 2 led us to define four cancer classes: Class 1 = [GBM], Class 2 = [SKCM], Class 3 = [PAAD] and Class 4 = [LUAD, LUSC, PRAD, STAD, HNSC, BLCA, LGG, THCA]. The seriation matrices in Figure 3a represent the binary classifier accuracies and sensitivity/specificity for these different classes.

**Figure 3.**
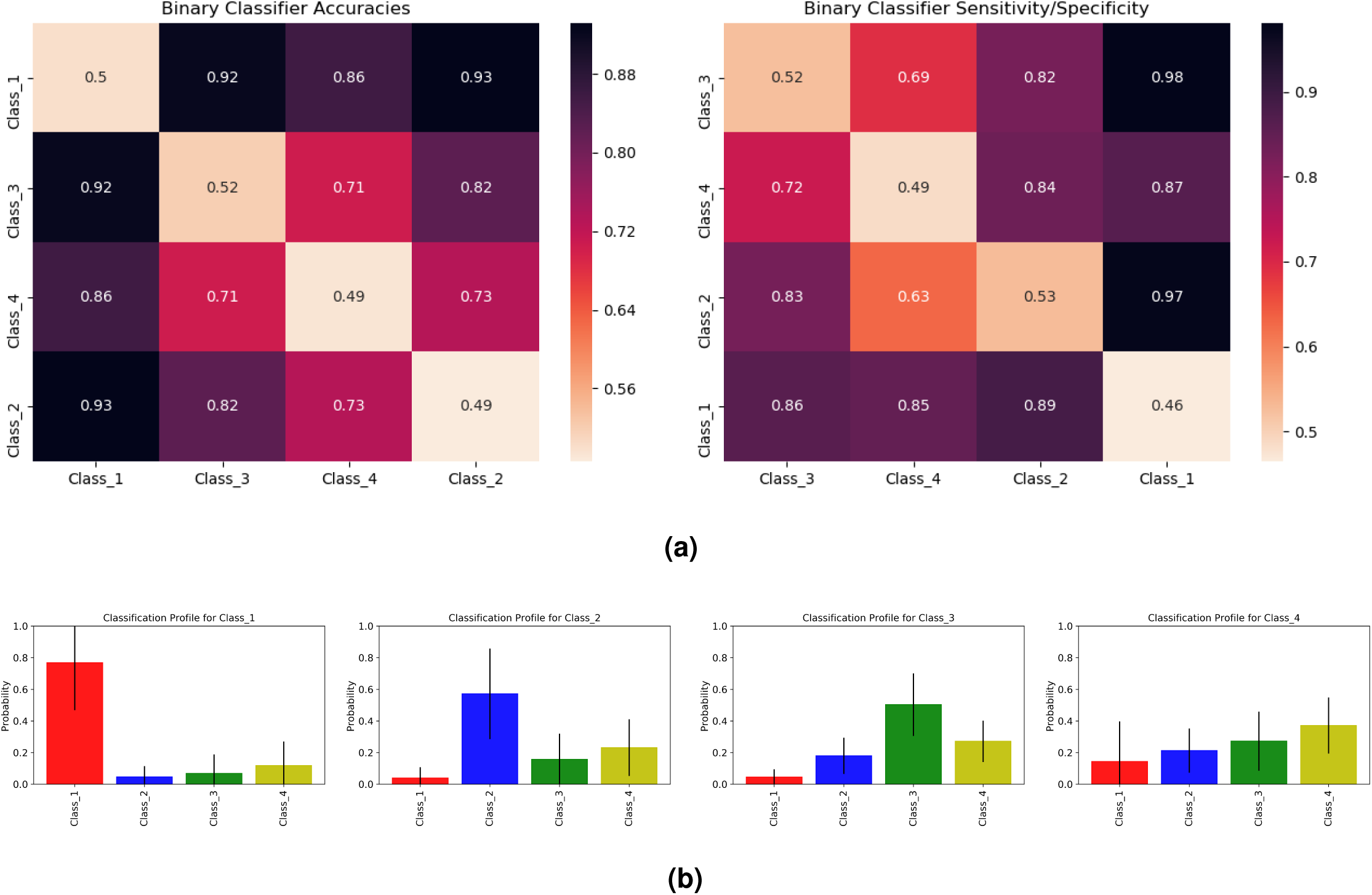
**(a)** These seriation matrices show 4-fold validation accuracy and sensitivity/specificity for the four main clusters of cancers in Figure 2 generated using 3843 blood-derived normal samples. Class 1 = (brain), Class 2 = (skin), Class 3 = (pancreas), Class 4 = (stomach, bladder, prostate, brain_lgg, lung, thyroid, lung_sq, head_neck). **(b)** Mean and standard deviations for the cancer classification profiles of individuals in Class 1, Class 2, Class 3 and Class 4 (viewing left to right). To generate these profiles, we a trained multiclassifier on all four classes of cancers using gradient boosting. We then used this multiclassifier to obtain cancer classification profiles for a different set of individuals reported the average results for each cancer class. Class 1 individuals show a high probability for Class 1 cancers. Class 2 and Class 3 individuals also show a higher probability for their respective classes, but with a slightly weaker signal. Class 4 individuals show similar probabilities for Class 2, Class 3 and Class 4.

### Cancer Classification Profiles

To assess a patient’s propensity of developing a class of cancers, we trained a multiclassifier for the four cancer classes using gradient boosting. This classifier uses a mutation profile to predict the relative probability of each class of cancer. Figure 3b shows the mean and standard deviation of these probabilities when tested on patients from each cancer class. Class 1, 2, and 3 all give large probabilities for their respective classes. Classes 2 and 3 give weaker signals because they are closer to Class 4 than Class 1 is (see Figure 3a). Individuals in Class 4 in the test set have similar scores for Classes 2, 3 and 4, showing that Class 4’s signal is not very distinct from Classes 2 and 3. This can again be attributed to the closeness of Class 4 to both Class 2 and Class 3 in the seriation diagram in Figure 3a. Supplementary Figure 6 gives the classification profile for Class 4 individuals when training only on Classes 1, 2 and 3 individuals. Again, Class 4 seems to imitate Classes 2 and 3, but the high standard deviation in Class 1 probability suggests Class 4 cancer patients can also have a high probability for Class 1 cancers.

### Effect of Adding NAT samples on classifiers

Recent studies have shown positive associations of Solid Tissue Normal (Normal Adjacent to Tumor (NAT)) samples on TCGA with the tumor DNA of cancer patients^25, 26^. We added 687 unamplified NAT samples as mentioned in Table 1A (Column 3) in our analysis to check if their presence is useful in discovering a stronger cancer-type signal. More precisely, we combined the 3874 blood-derived and 687 NAT samples to construct the pairwise classifiers. Here, we also covered TCGA-KIRC as now we had 210 (179 NAT and 31 blood-derived) samples that were enough to build reliable classifiers. We didn’t observe any significant improvement in cancer signal detection by adding NAT samples over only using blood-derived normal samples and found the same cancer classes that we discovered previously (see Figure 4a, Supplementary Figure 7). Further, we found that TCGA-KIRC belonged to the same class as TCGA-GBM showing strongly distinctive signal from the other 10 cancers (see Figure 4a, Supplementary Figure 7).

**Figure 4.**
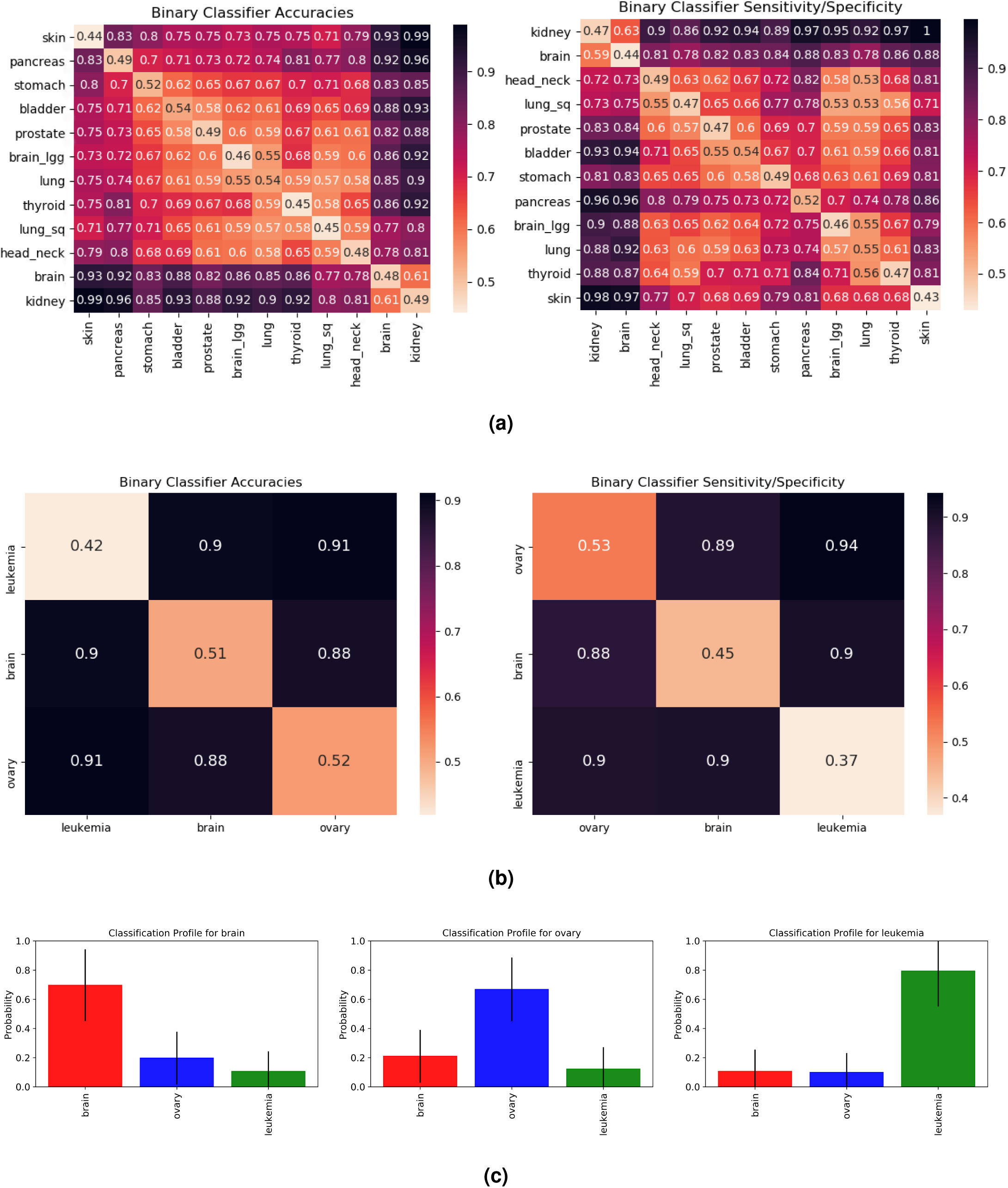
**(a)** The matrix on the left contains the pairwise validation accuracies of classifiers built by using 4561 unamplified samples covering both blood-derived and solid tissue normal DNA of 12 different cancer types. These cancer types are TCGA-SKCM (skin), PAAD (pancreas), STAD (stomach), BLCA (bladder), PRAD (prostate), LGG (brain_lgg), LUAD (lung), THCA (thyroid), LUSC (lung_sq), HNSC (head_neck), GBM (brain), KIRC (kidney). **(b)** Classifier accuracies and sensitivity/specificity when training and testing is done using amplified samples only. The cancers covered here are TCGA-GBM (brain), TCGA-OV (ovary) and TCGA-LAML (leukemia). There is a strong signal distinguishing all these cancer types. We speculate that TCGA-KIRC would behave similarly to TCGA-GBM, since these cancers are similar in Figure 4a. **(c)** Mean and standard deviation for the cancer classification profiles of individuals belonging to TCGA-GBM (brain), TCGA-OV (ovary) and TCGA-LAML (leukemia). These classification profiles are obtained by building a multiclassifier using gradient boosting. The multiclassifier is obtained by training *only* on amplified healthy samples of brain, ovary and leukemia cancers on TCGA. The testing is done on a separate set of amplified samples for these cancers. It can be seen that brain, ovary and leukemia cancer patients in the test set are showing a higher probability for the respective cancer using their healthy DNA mutation profile.

### Analysis of Amplified Samples

Amplification techniques have been shown to bias tandem repeat information^27^, especially in TCGA data^28^. To control for this, we separately analyzed amplified samples. Figure 4b shows the seriation diagrams for accuracy and sensitivity/specificity of pairwise classifiers built using 525 samples *amplified* by MDA technology. Because of the limited data, this test only covered TCGA-GBM (brain), TCGA-OV (ovary) and TCGA-LAML (leukemia) and both the normal DNA types, i.e. blood-derived and solid tissue normal (NAT) were used (Table 1B, Supplementary Files 13-15). The high accuracy and sensitivity/specificity values in these diagrams suggest a strong cancer-type signal in the mutation profiles of the healthy DNA. Further, we also generated the classification profiles using these amplified samples for individuals with brain, ovary and leukemia cancer. Figure 4c shows the mean and standard deviation of the predicted cancer probabilities for these three populations. The highest probability cancers correspond with the cancers that the patients were diagnosed with, affirming that healthy DNA contains a cancer-type signal.

Genome analysis for the results presented in Figures 2-5 and Supplementary Figures 6-14 was done using samtools^29^ (see Methods, Code/Software) and the pipeline presented in Figure 1b. We also verified these results for unamplified samples by using another genome analysis tool for short tandem repeats (STR)-hipSTR^30^ that only detects tandem repeats with pattern lengths atmost 6 (see Supplementary Figure 15).

### Driver Genes

Studies in the past have identified driver genes like TP53, BRCA-1, BRCA-2, etc. We considered 723 such genes that are listed in Supplementary File 16 obtained from Cancer gene census - COSMIC^31, 32^.

To test whether these regions provided special information, we filtered our mutation profiles to only use tandem repeats that overlapped with driver gene regions. We conducted this experiment for the 4561 unamplified samples and 525 amplified samples separately. Figure 5 shows a comparison the classifiers trained on these filtered mutation profiles and mutation profiles which contain all the features *except* those in the filtered profiles. Darker cells in this figure correspond to large differences in the accuracy of the classifiers, indicating that these signals exist outside of driver gene regions. We notice these darker cells especially for TCGA-PAAD (pancreas) and TCGA-SKCM (skin). We also see noticeable differences when TCGA-OV (ovary) is compared against TCGA-GBM (brain) and TCGA-LKCM (leukemia). The driver-gene classifiers always performed worse than the classifiers trained on the rest of the genome, indicating that the signal exists both inside and outside driver gene regions.

**Figure 5.**
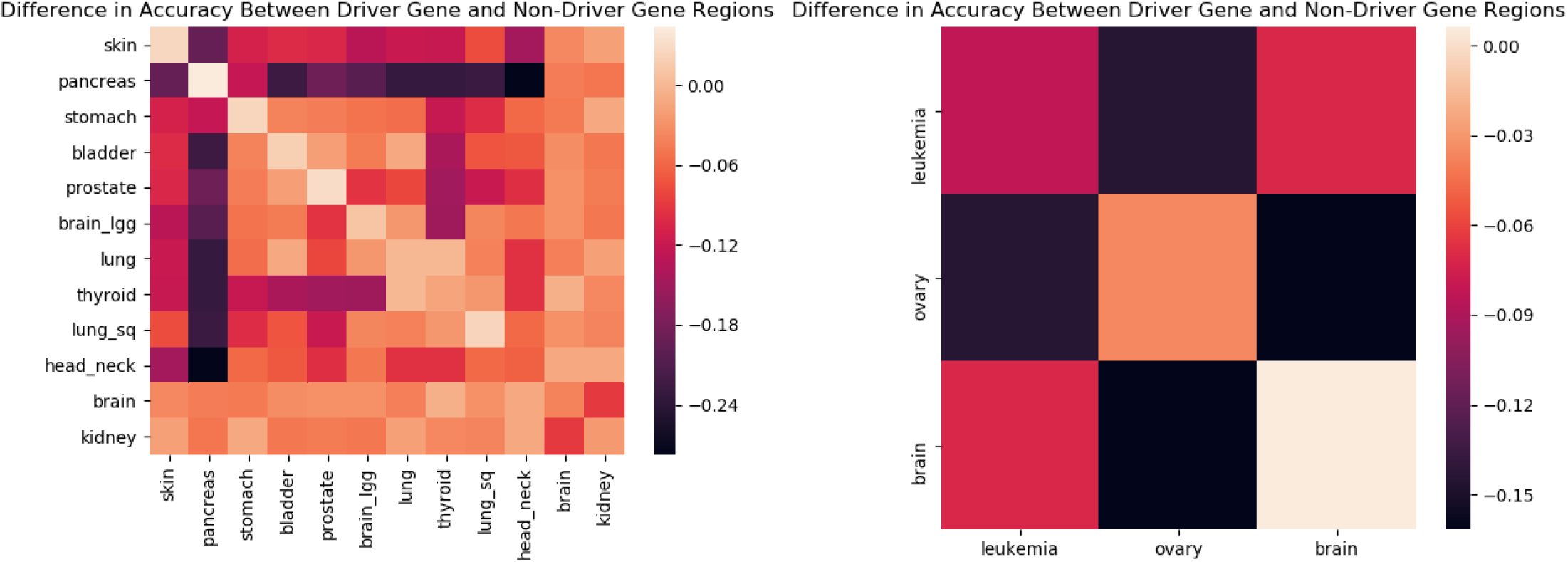
A comparison of classifiers trained on mutation profiles restricted to known driver-gene regions and those restricted to non-driver-gene regions. Darker cells represent a larger difference in the testing accuracies of the classifier, indicated by the scale. This experiment was performed on unamplified (left) and amplified (right) samples separately. The uneven coloring suggests that some cancer-type signals exist more primarily in driver-gene regions than others.

## Discussion

### Early Cancer prediction

We have shown that the mutation profiles of the blood-derived normal genomes of cancer patients contain a cancer-type signal (Figures 2 & 3). It is reasonable to assume that the mutation profiles of cancer-free patients may also contain these signals, and we can use our classifier to quantify their presence. The cancer classification profiles given by this classifier could be used to screen individuals for those who may benefit from more comprehensive and expensive cancer detection tests.

### Accumulation of Mutations

Searching for information-containing features within 3 billion nucleotides is a formidable task. This has traditionally been simplified by comparing individuals to extract variants, which compresses the genome into a smaller set of features to analyze. These differences, known as SNPs and CNVs, are central to both Mendelian studies^5^ and GWAS^6, 7, 12^.

This form of genome compression loses crucial information about how the genome is *changing* by only considering differences in the genome’s *current state*. Every individual’s genome passes through a distinct evolution channel that is controlled by hereditary, environmental and stochastic factors. These evolution channels differ among the population and can give rise to different risks of disease, but we cannot easily identify these differences from the single-generation SNP and CNV analysis used in GWAS. Mendelian studies may provide insight into inter-generational processes, but do so at the cost of requiring inter-generational data, which severely limits the scope of a feature search. Even with additional data, Mendelian studies still lack the ability to detect differences in mutation processes that occur throughout one’s lifetime.

Mutation profiles are generated without any comparative analysis, reducing the data-demand. Instead, the tandem repeat regions in a *single* genome provide a window into its history, capturing information about the individual’s evolution channel. This ability to reconstruct a genome’s history from repeat regions is lost when studies only view differences between individuals. The use of mutation profiles expands our access to time-dependent traits which may be essential to understanding developed diseases like cancer.

### Sequencing Technology Limitations

TCGA samples are obtained from Illumina platforms with a coverage depth 30-40X. The read lengths used are short ranging between 100-500 bp. This poses a problem in the detection of longer tandem repeats^33–35^. In our analysis, we only used repeats with pattern lengths ≤ 10 bp and number of copies not greater than 100.

## Methods

### WXS data

We used exome data from “blood derived normal” and “solid tissue normal” samples in the TCGA^20^ database, details about which are provided in the Supplementary Files 1-15. The BAM file for each sample was aligned against hg38. All the autosomes from each sample were recovered using samtools^29^.

### Algorithms

Our algorithms are partitioned to Part A and Part B (see Fig. 1b). Part A is only performed once, where Part B is performed whenever cancer prediction is required. In Part A, a dataset of healthy DNA is first processed by the Benson^21^ and Tang *et al*.^22^ algorithms to deduce the mutation profiles. Then, these vectors are aligned by a dynamic programming algorithm to resolve missing regions. Finally, the aligned vectors are fed into a training algorithm to produce a classifier. In Part B, this classifier is applied over any individual’s genome, to assess the overall probability to contract any of the cancer in question.

### Tandem Repeat Detection and Duplication History Estimation

*Tandem duplications* are consecutively repeated patterns caused by replication slippage events^36, 37^, in which a pattern is duplicated next to the original. For example, the following shows two tandem duplications of length 4, where the duplicated part is highlighted in bold. The underlined segment is the *microsatellite* or *repeat region*.

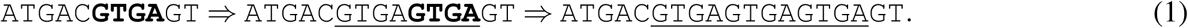

The *pattern* of a region is the short strand which repeats itself. The *copy number d* of a repeat region indicates the number of times that the pattern is repeated. For example, the pattern of the underlined repeat region in the right hand side of (1) is `GTGA`, and its copy number is 3.

Microsatellites are usually accompanied by various types of errors: substitutions (replacement of one nucleotide by another), deletions (omission of a nucleotide), and insertions (addition of a nucleotide). The total number of substitutions, deletions, and insertions in a repeat region is called the *error number m*. For example, the following shows the contamination of (1) by 1 substitution, 1 deletion, and 1 insertion.

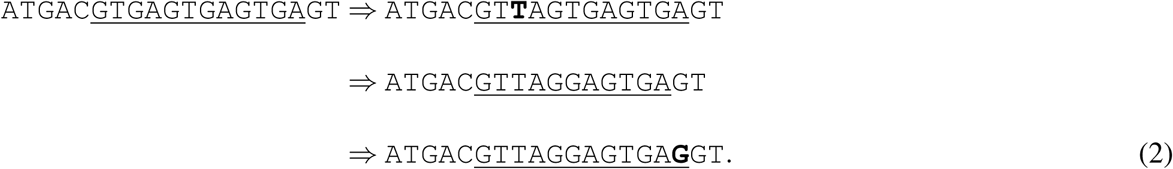

Clearly, the copy number of (2) is 3 and its error number is 3, and hence its mutation index is (*m, d*) = (3, 3). In the first step of Part A we use the Benson Tandem Repeat Finder to detect repeats with consensus pattern size at most 10 and copy number at most 100. These size limitations mean we only consider regions smaller than 1000 nucleotides. The single block version of the duplication history estimation algorithm given in Tang *et al*.^22^ was then applied to each tandem repeat region to obtain the respective *mutation index* = (*m, d*). The aggregation of these (*m, d*) values gives a vector twice the size of the number of repeat regions, which we call an individual’s *mutation profile*. Since, TCGA data is WXS, we only calculated a unique *mutation profile* of an individual’s exome.

### Alignment

Following the completion of the Benson and Tang *et al*. algorithms, it was sometimes the case that certain repeat regions appeared in some patients and did not appear in others. In addition, minor differences were observed in the patterns of identical repeat regions in different individuals. As a result, a technical difficulty arose in handling the input to the learning algorithm. Consider the following two patients, in which the repeat regions are underlined.

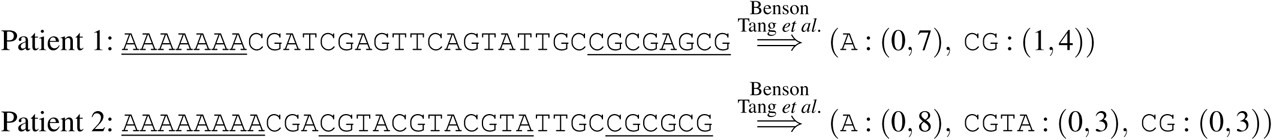

The success of machine learning depend on the detection of patterns in specific *positions* of feature vector, so entries which correspond to the same repeat region must also be placed in the same position for all inputs. This is clearly not the case in the above example, in which the second entries of the vectors correspond to different repeat regions.

This issue is resolved by using a dynamic programming alignment algorithm. In this algorithm, a similarity score is computed recursively for each possible alignment, and the alignment which leads to the best possible score is chosen. Each possible alignment is defined as the sum of normalized *edit-distances*^1^ between the patterns of all respective pairs. Further, the distance between any pattern and a “missing pattern”, denoted by ‘–’ below, is defined as 0.4. Namely, two patterns whose respective normalized edit distance is less than 0.4 were considered to be equal for the sake of the alignment. For example, the vectors above are aligned in the following way.

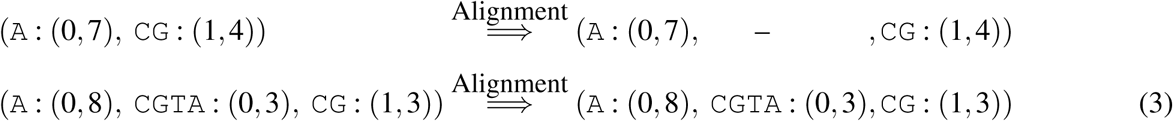

The score for the alignment (3) is *d*_*e*_(`A`, `A`) + *d*_*e*_(–, `CGTA`) + *d*_*e*_(`CG`, `CG`) = 0 + 0.4 + 0 = 0.4, where *d*_*e*_ denotes edit distance. For comparison, the alternative alignment

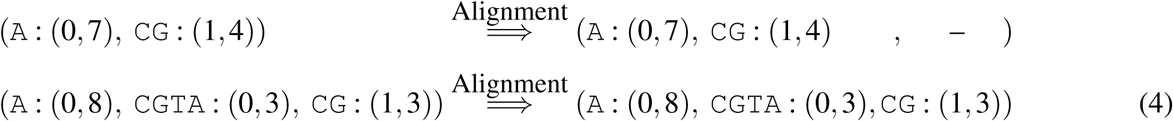

has score of *d*_*e*_(`A`, `A`) + *d*_*e*_(`CG`, `CGTA`) + *d*_*e*_(–, `CG`) = 0 + 2*/*3 + 0.4 ≈ 1.06, and hence (3) is preferred over (4).

The mutation profile of each individual was aligned against the mutation profile of the reference genome (hg38) by using the method that is mentioned above. The repeat regions that were missing in the reference genome were omitted from these aligned mutation profiles. Further, given the aligned mutation profiles, every ‘–’ is replaced by (0, 0). This gave aligned mutation profiles of the same size that can now be used as features for the learning part described next.

### Machine Learning

The aligned mutation profiles were used as features for the learning algorithm. Machine learning classifiers for distinguishing cancers were obtained using two approaches:

#### Pairwise Classifiers

We trained a binary classifier for *every pair* of types of cancer, generating 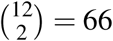 pairwise classifiers for unamplified samples and 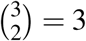 pairwise classifiers for amplified samples. The accuracy in either of those classifiers is used as a measure for the “uniqueness” of the mutation profiles that cause a certain type of cancer, and can additionally be seen as a distance measure between different types of cancer. We used xgboost^23^ algorithm at default parameters with max-depth = 2, and performed 4-fold validation to build each of these pairwise classifiers.

#### Multiclassifier

This was built using xgboost ‘multi:softprob’ parameter with max-depth = 2 and predicted the probability of all the cancers simultaneously. Again 4-fold cross validation was performed to avoid over-fitting.

### Code/Software

The code and necessary documentation for the pipeline used is available at http://paradise.caltech.edu/~sidjain/Codes.tar.gz.

### Data Availability

The BAM files for WXS samples of cancer patients used in the study were obtain from The Cancer Genome Atlas (TCGA)^20^. These files have controlled access and cannot be availed publicly. However, request to access TCGA controlled data can be made via dbGap^38^ (accession code: phs000178.v1.p1). The file names for the analyzed samples are given in Supplementary Files 1-15.

## Supporting information

LUSC

PAAD

LUAD

HNSC

SKCM

KIRC

GBM

PRAD

STAD

THCA

LGG

BLCA

GBM_W

LAML

## Acknowledgements

This work was supported in part by The Caltech Mead New Adventure Fund and a Caltech CI2 Fund. The authors would like to thank Eytan Ruppin for his valuable advice and feedback.

## Author contributions statement

S.J. analyzed the TCGA genomic data, implemented repeat finding and history estimation steps in the pipeline, helped with the machine learning step in building pairwise and multiclassifiers, and wrote the manuscript; B.M. implemented the machine learning pipeline; N.R. implemented the alignment algorithm; J.B. originated and guided the study. S.J., B.M., N.R. and J.B. participated in brainstorming of the concepts and discussions and revisions of the manuscript.

## Ethics Statement

The ethics approval to the TCGA data was granted by Caltech Institutional Review Board.

**Figure 6. (Supplementary).**
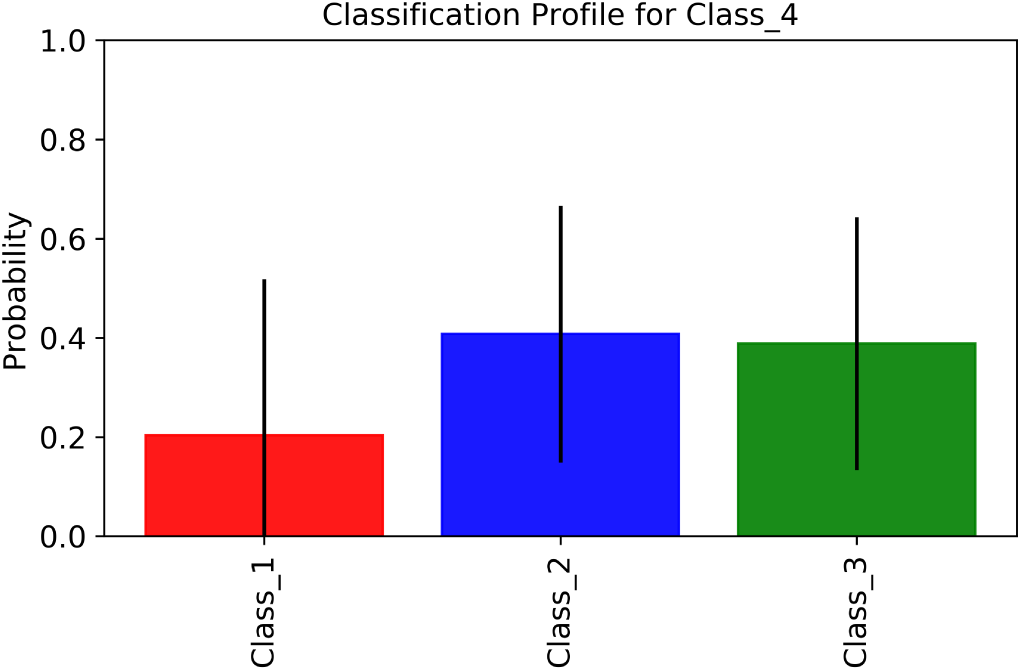
Cancer classification profile for Class 4 individuals when the multiclassifier was trained only using Class 1, Class 2 and Class 3 samples mentioned in Figure 3a. This shows a stronger association of Class 4 with Class 2 and Class 3 than Class 1. However the high standard deviation for Class 1 also means that some individuals in Class 4 have a stronger association with Class 1 than Class 2 and Class 3.

## Supplementary Information

We replicated our experiments with only error number *m* and only copy number *d* values in the mutation profiles. These experiments gave results similar to the complete profiles, suggesting that both *m* and *d* contain cancer-type signals. Supplementary Figure 9 shows the associated pairwise accuracies and sensitivity/specificity for cancers with 3843 blood-derived normal unamplified samples when only *d* was used in the mutation profile and Supplementary Figure 10 shows the associated pairwise accuracies and sensitivity/specificity when only *m* was used. Supplementary Figures 11 and 12 show the plots using all the 4561 unamplified samples. Supplementary Figures 13 and 14 show the plots for the 525 amplified samples..

**Figure 7. (Supplementary).**
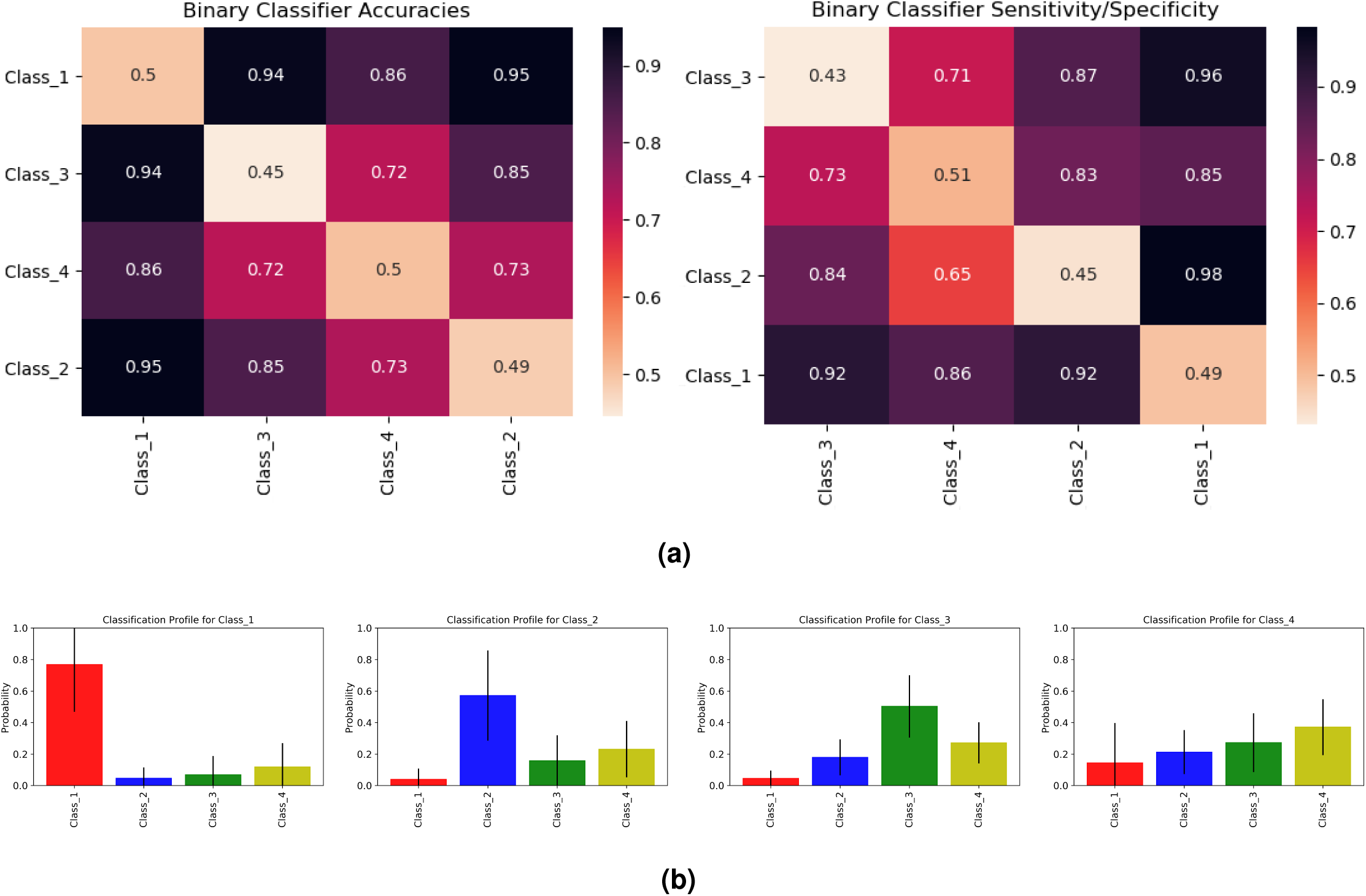
**(a)** These seriation matrices show 4-fold validation accuracy and sensitivity/specificity for the four main clusters of cancers in Figure 4a where all the 4561 unamplified samples were used. Class 1 = (brain, kidney), Class 2 = (skin), Class 3 = (pancreas), Class 4 = (stomach, bladder, prostate, brain_lgg, lung, thyroid, lung_sq, head_neck). **(b)** Mean and standard deviations for the cancer classification profiles of individuals in Class 1, Class 2, Class 3 and Class 4 (viewing left to right). To generate these profiles, we a trained multi-classifier on all four classes of cancers using gradient boosting. We then used this multi-classifier to obtain cancer classification profiles for a different set of individuals reported the average results for each cancer class. Class 1 individuals show a high probability for Class 1 cancers. Class 2 and Class 3 individuals also show a higher probability for their respective classes, but with a slightly weaker signal. Class 4 individuals show similar probabilities for Class 2, Class 3 and Class 4.

**Figure 8. (Supplementary).**
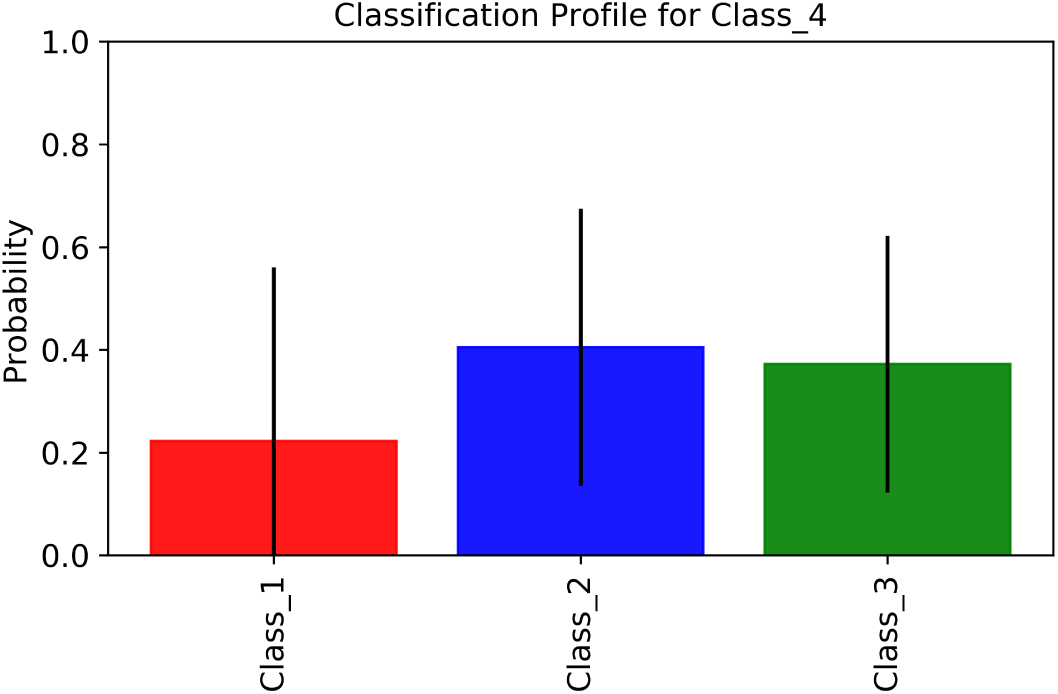
Cancer classification profile for Class 4 individuals when the multiclassifier was trained only using Class 1, Class 2 and Class 3 samples shown in Supplementary Figure 7. This shows a stronger association of Class 4 with Class 2 and Class 3 than Class 1. However the high standard deviation for Class 1 also means that some individuals in Class 4 have a stronger association with Class 1 than Class 2 and Class 3.

**Figure 9. (Supplementary).**
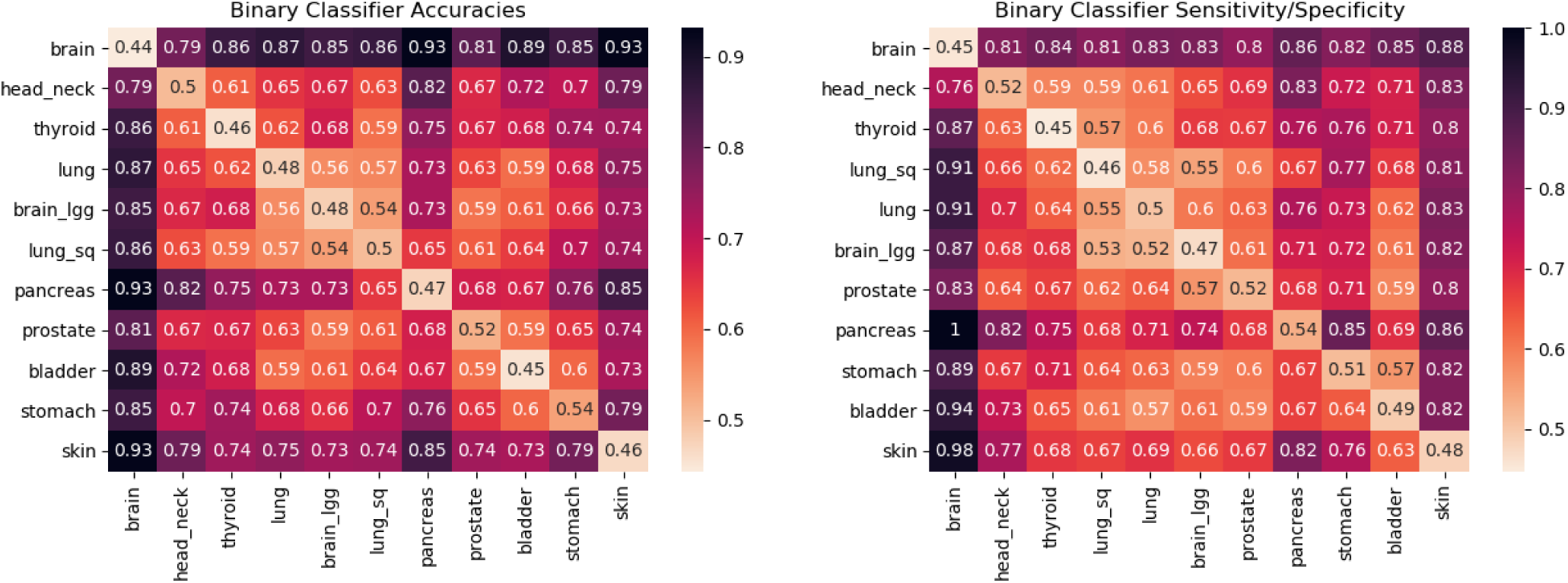
Accuracies and sensitivity/specificity for the pairwise classifiers when only the copy number (*d*) information was used in the mutation profile for the 3843 WXS blood-derived unamplified samples mentioned in Table 1A. Notice that like Figure 2, we again find strong distinguishing signals for brain, pancreas and skin cancers as can be seen by the darker rows corresponding to these cancers.

**Figure 10. (Supplementary).**
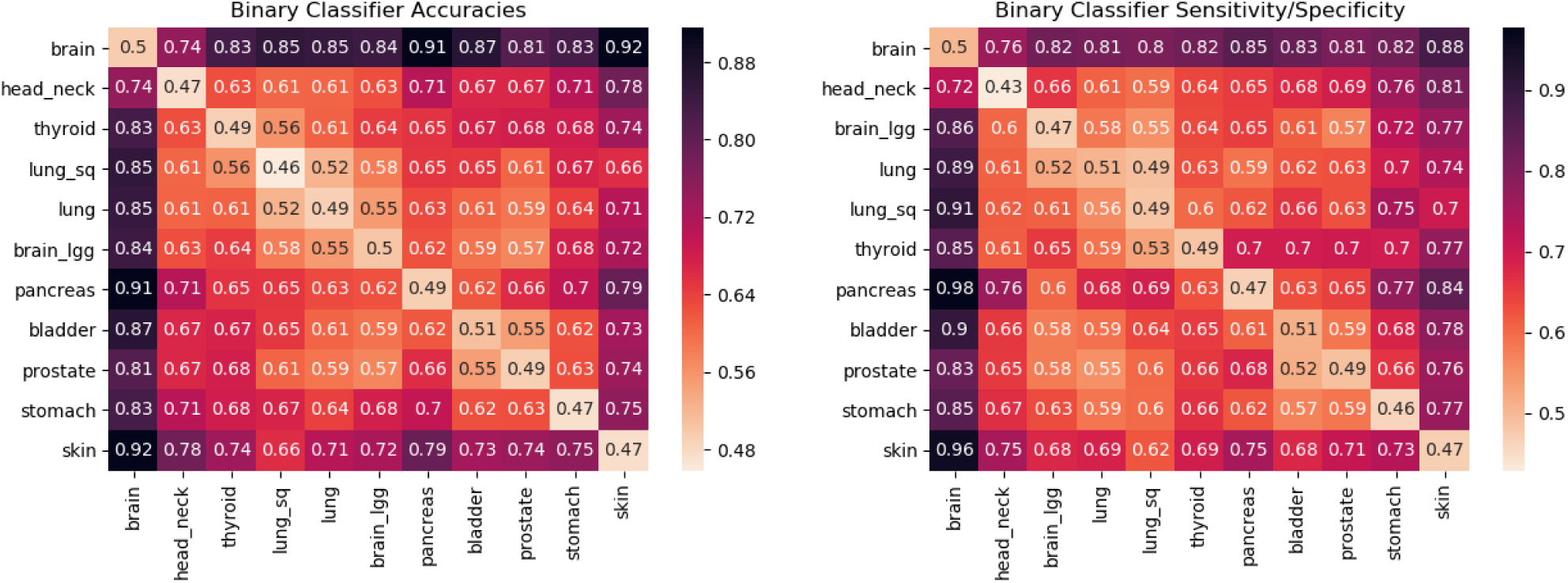
Accuracies and sensitivity/specificity for the pairwise classifiers when only the error number (*m*) information was used in the mutation profile for the 3843 WXS blood-derived unamplified samples mentioned in Table 1A. Notice that like Figure 2, we again find strong distinguishing signals for brain and skin cancers as can be seen by the darker rows corresponding to these cancers. However unlike Figure 2, the signal for pancreas cancer is not as strong.

**Figure 11. (Supplementary).**
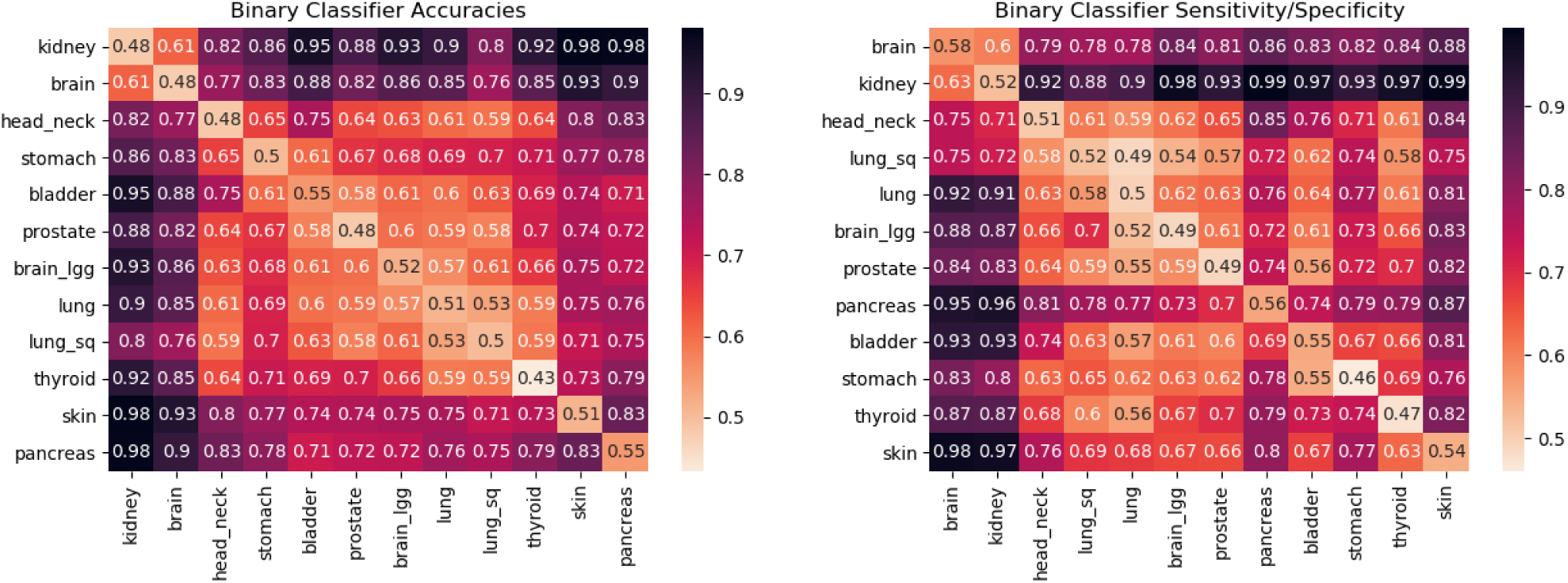
Accuracies and sensitivity/specificity for the pairwise classifiers when only the copy number (*d*) information was used in the mutation profile for the 4561 WXS unamplified samples that comprised both blood-derived and solid tissue normal type DNA as mentioned in Table 1A.Notice that like Figure 4a, we again find strong distinguishing signals for (brain,kidney), pancreas and skin cancers as can be seen by the darker rows corresponding to these cancers.

**Figure 12. (Supplementary).**
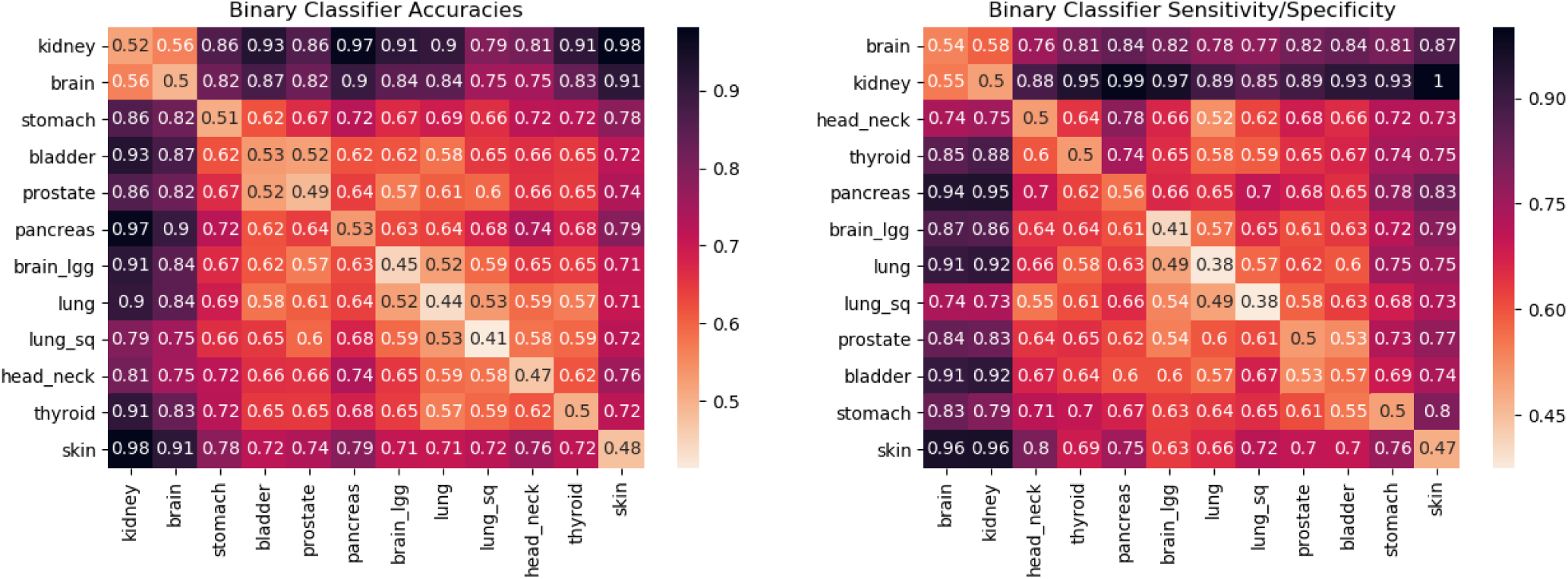
Accuracies and sensitivity/specificity for the pairwise classifiers when only the error number (*m*) information was used in the mutation profile for the 4561 WXS unamplified samples that comprised both blood-derived and solid tissue normal type DNA as mentioned in Table 1A.Notice that like Figure 4a, we again find strong distinguishing signals for (brain,kidney) and skin cancers as can be seen by the darker rows corresponding to these cancers. However, unlike Figure 4a, the signal for pancreas cancer is not as strong.

**Figure 13. (Supplementary).**
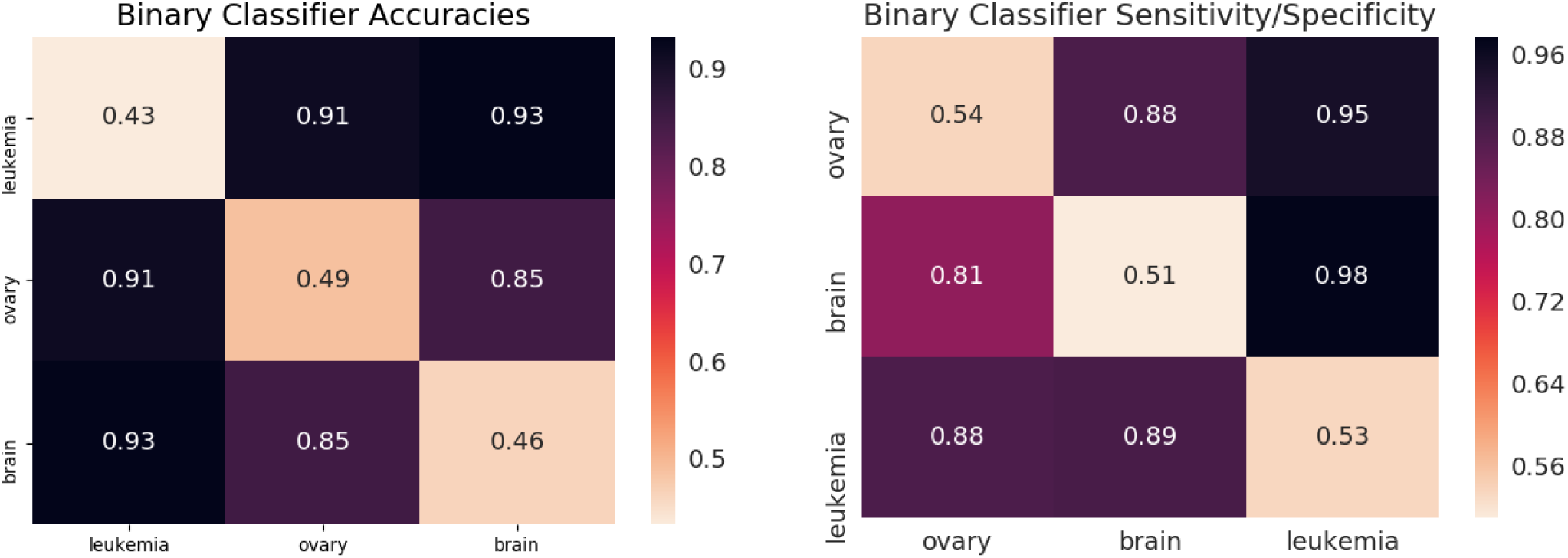
Accuracies and sensitivity/specificity for the pairwise classifiers when only the copy number(*d*) information was used in the mutation profile for the 525 WXS amplified samples mentioned in Table 1B. Notice that like Figure 4b, we again find strong distinguishing signals for all the three cancers.

**Figure 14. (Supplementary).**
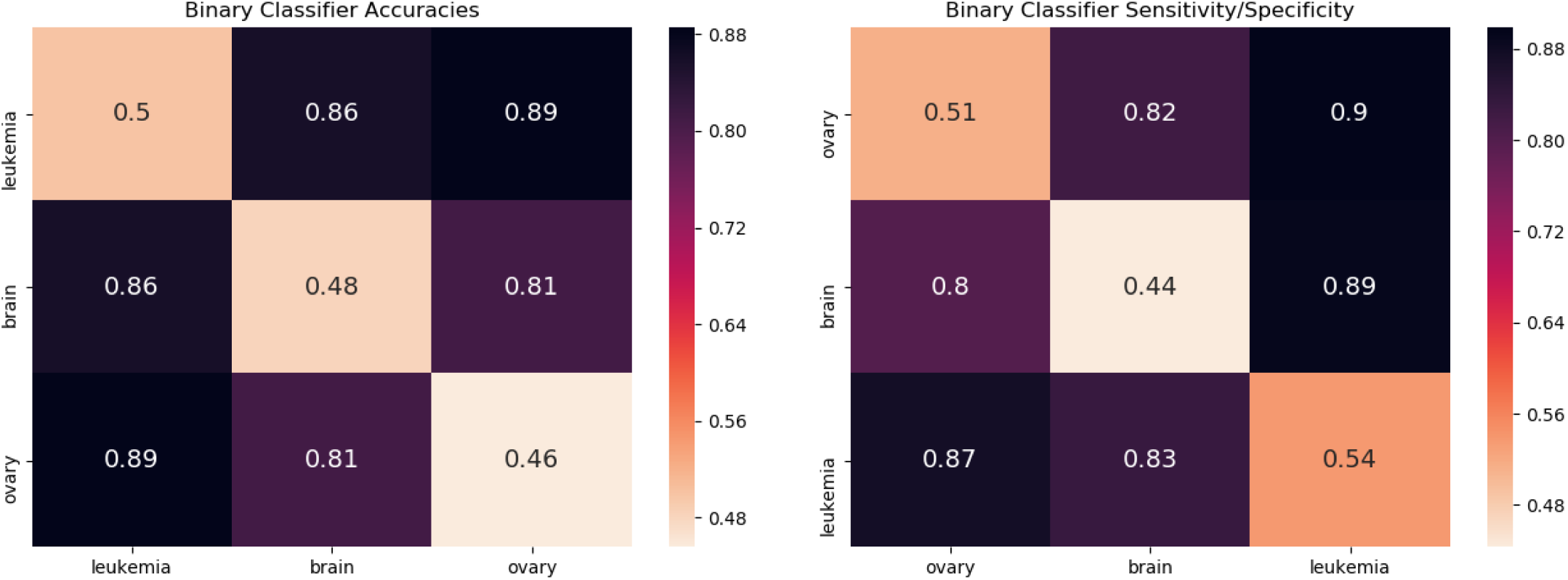
Accuracies and sensitivity/specificity for the pairwise classifiers when only the error number (*m*) information was used in the mutation profile for the 525 WXS amplified samples mentioned in Table 1B. Notice that like Figure 4b, we again find strong distinguishing signals for all the three cancers.

**Figure 15. (Supplementary).**
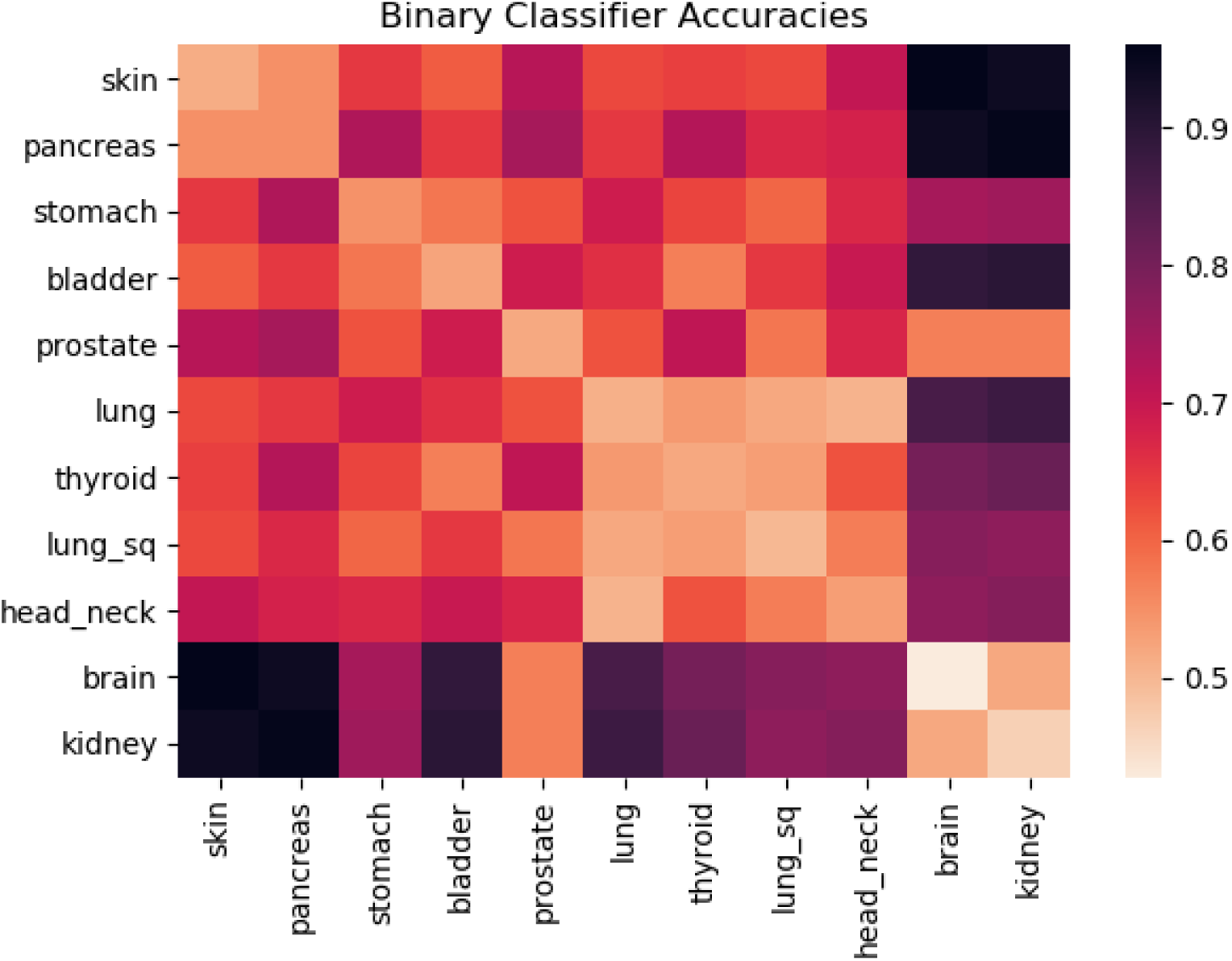
Accuracies for the pairwise classifiers when hipSTR was used for STR detection. hipSTR detects short tandem repeats with pattern lengths 6 or less. We used **–min-reads** 1 **–def-stutter-model** setting. Even though hipSTR requires further trimming of mutation profiles to tandem repeats with pattern lengths ≤ 6, we were still able to detect cancer-specific signals. The differences from Figure 4a in Figure 15 (for example: prostate and brain, prostate and kidney, skin and pancreas) can be attributed to the fact that there are tandem repeat regions with pattern lengths greater than 6 that contained critical cancer-specific information. Further, we only used STRs that were commonly detected by hipSTR in all the samples being analyzed. Therefore, the STRs that were not detected in some samples were not used in this analysis.

## Footnotes

1 That is, the minimal number of insertions, deletions, and substitutions that are required to transform one pattern to the other, divided by the average length of the sequences.

